# Exon shuffling hotspots in an animal genome

**DOI:** 10.1101/2025.10.28.685065

**Authors:** Caroline M. Weisman

## Abstract

The birth of new genes is key to evolution. Exon shuffling is a powerful mechanism for gene birth in which new genes are made by stitching together fragments of existing ones. I report hotspots of exon shuffling in the genome of the animal *Drosophila affinis*. These are relatively small regions that quickly receive, concentrate, and rearrange gene fragments from around the genome. I detail the recent creation of transcripts containing composite open reading frames at these hotspots, including cases that resemble not just the birth of a single gene but of gene families. I describe basic phenomenology of the hotspots and propose that they are best described not specifically in terms of exon shuffling but more broadly by their tendency to toy with existing genomic sequences as “tinkering loci.” I speculate that tinkering loci may overcome barriers to the reassortment of modular sequences, accelerating animal evolution.

## Introduction

The gene is the atom of biology: the basic functional unit from which biological structures are built. Understanding their birth is essential for a meaningful understanding of evolution.

One type of mechanism generates new genes by reassorting pieces of old ones. This is a powerful strategy: there is novelty to be had in combining existing functions in new ways, and it is easier for evolution to re-use what it has already invented.

“Exon shuffling” is one such mechanism. It occurs when recombination between different regions of the genome physically mixes fragments of different genes, creating a new gene that is their composite [1]. Introns enable exon shuffling by increasing the target size for recombination breakpoints that can transfer genic sequence. Repetitive sequences enable exon shuffling by providing similar sequences at different loci, which is required for ectopic recombination [2][3].

Exon shuffling is very rare in most taxa, but feasible in animal genomes, which have large introns and many transposon-derived repeats. Exon shuffling seems to have created many of the gene families on which animal multicellularity is based, including cell adhesion molecules, cell-cell signaling receptors, and proteins that make and form extracellular matrices [4].

Even in animal genomes, there are at least two barriers to gene birth by exon shuffling. First, in a large animal genome, it is unlikely that independently shuffled fragments will be moved close enough to one another. Second, it is unlikely that they will be moved into the right relative orientation – including frame, order, and spacing – to make a functional protein.

Combining pieces of genes that are already neighbors can happen despite the first barrier and is the most common form of exon shuffling [3,5]. The second barrier can be mitigated by limiting shuffling to introns in the same frame, which seems to have been key in the production of the animal multicellularity toolkit mentioned above [6]. These workarounds substantially limit the combinations of gene fragments available for the formation of new genes.

## Results

### An exon shuffling hotspot in the *Drosophila affinis* genome

I came across an unusual region in the genome of *Drosophila affinis*, a fruit fly that diverged from *Drosophila melanogaster* about 30 million years ago [7]. This region appears to be a “hotspot” for exon shuffling (Figure 1).

**Figure 1:**
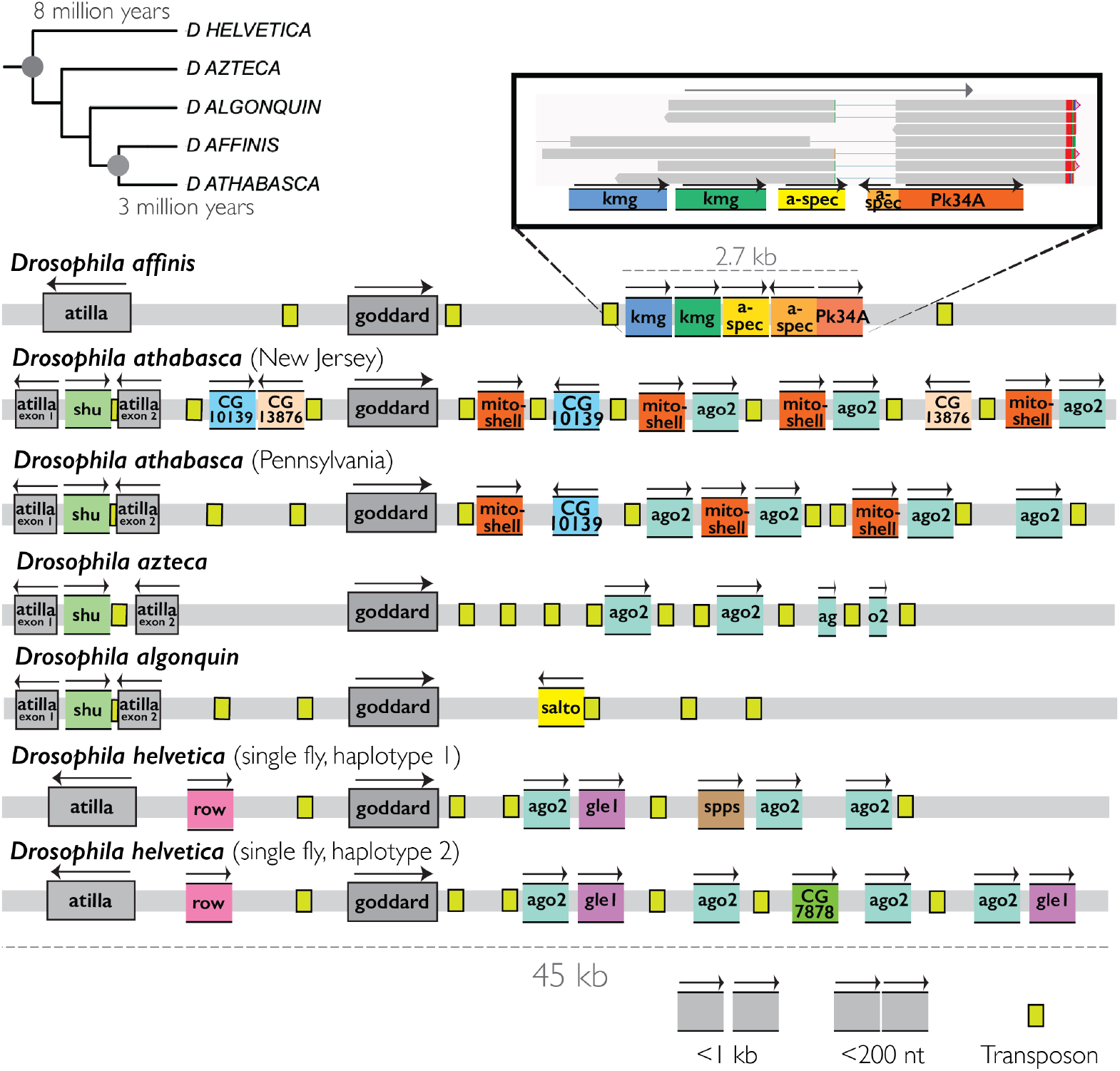
An exon shuffling hotspot. The same genomic region in *Drosophila affinis* and six genome assemblies from four closely related (<8 million years) species. Phylogeny and times of key nodes shown in the upper left. Gray bars are chromosomes; bordered rectangles are genes that are full-length relative to the *melanogaster* copy, used as a proxy for the ancestor; truncated rectangles with no side borders are fragments of genes relative to the *melanogaster* copy. Gray genes are those whose locations are conserved in the ancestor of this group; colors indicate genes or fragments that are derived. Each unique color denotes fragments that are subsets of one another or that have very similar boundaries: for example, all *argonaute2 (*ago2, turquoise) fragments cover varying intervals of the same 250 amino acid region, overlapping its piwi domain. Gene names are the FlyBase [8] (www.flybase.org) gene symbol for the most significant *melanogaster* homolog. For legibility, spacing is **<u>not to scale</u>**, but effort has been taken to give an accurate qualitative sense, and two important classes of distances between closely spaced elements, as indicated in the bottom right, are depicted. The same region shows significant turnover, duplication, and repositioning of gene fragments between these closely related species. Inset in the upper right shows an IGV [9] visualization of long-read “Iso-seq” RNA sequencing data of reads from a complex of fragments in *Drosophila affinis*. Horizontal gray bars are single transcripts; thin lines are spliced introns; red or green mismatches at the end of the transcripts are the polyA (red, A, plus strand; green, T, minus strand) tails. Five fragments from three genes are expressed together on the same transcript. The splice boundaries correspond roughly to the edges of two of the fragments.

In five closely related species spanning about 8 million years of divergence [7], the locus has a high density of gene fragments, derived from many different genes in different regions of the genome. The sequences surrounding many of the fragments match intergenic and intronic sequence surrounding their parent genes, suggesting ectopic recombination as the mutational mechanism. Other fragments correspond to internal segments of exons and have no flanking sequence to support recombination, but are still consistent with it: recombination breakpoints could have been within the exon, or the flanking sequence could have been degraded. The alternative of retrotransposition requires the degradation of almost the entire transcript on the short timescales here, making it seem unlikely to be a major source.

The identity of the fragments varies substantially between species. Each species has a different assemblage, and most fragments are species-specific. There are also differences between different populations of the same species (*athabasca* [10]), and, based on a phased assembly, the two chromosomes of an individual fly (*helvetica* [11]). On these shorter timescales, many structural mutations within the locus itself, like the duplication of fragments that are already present, or their rearrangement relative to one another, can be resolved.

In all species, the region is rich in transposable elements. These may provide the sequence similarity required for both ectopic recombination with other loci and internal rearrangements.

To determine whether these products of shuffling are expressed, I acquired the strain of *Drosophila affinis* from which the assembly used here [12] was generated and performed Pacbio long-read “Iso-seq” RNA sequencing (Methods), in which a single read covers a whole transcript, on whole male flies. This revealed that a complex of five gene fragments at this locus, made from noncontiguous pieces of three different genes, is expressed together on the same transcript, expressed at 2 CPM (Figure 1, inset).

Available short-read RNA sequence data from sister species *athabasca* [13] showed that the fragments at this locus, despite none being in common with *affinis*, are also expressed, though it is not clear whether they are on the same transcript.

Intrigued by this hotspot, I searched systematically for more. In brief (Methods), I developed a simple custom genome annotation pipeline that identifies and retains small gene fragments, which are often discarded. I also annotated transposon sequences. I independently annotated available genome assemblies of *Drosophila affinis* and seven outgroup species (Resources and Data Availability), including four closely related species comprising the “*affinis* subgroup” that share a common ancestor only ∼8 million years ago. I mapped the Iso-seq data from *Drosophila affinis* to the *affinis* genome and used my annotation to identify “composite transcripts”: those containing coding fragments derived from two or more ancestral genes. I then used my annotations to determine whether the corresponding locus, with these fragments intact, was present in each of the other species. This allowed me to identify new composite transcripts and estimate their age.

### Hotspots create young composite open reading frames

Coding fragments on the same transcript will not produce a composite protein unless they form an intact open reading frame (ORF). The composite *affinis* transcript above (Figure 1) does not encode such an ORF. I was curious whether these hotspots have successfully produced such ORFs, and if so on what timescales.

I looked for composite ORFs that are present in *Drosophila affinis* and absent in two close (*Drosophila pseudoobscura, Drosophila obscura*) and one more distant (*Drosophila melanogaster*) outgroup species to the *affinis* subgroup, consistent with having evolved within the group in the last 8 million years. I found nine such composite ORFs, eight of which are specific to *Drosophila affinis* and thus younger than ∼3 million years. These are expressed at an average of 5 CPM, but with wide variance: four at <1 CPM (1-5 reads), three at 2-3 CPM, and one at 30 CPM.

Because they more clearly display the generative rate and underlying mechanisms of the *affinis* hotspots, I focus on these youngest composite ORFs. Below I detail three cases.

#### Mastermind/six-banded

One *affinis-*specific composite ORF was formed at an 11 kb locus that has gained 10 fragments from 10 different genes since the divergence of *affinis* from all other species (Figure 2).

**Figure 2:**
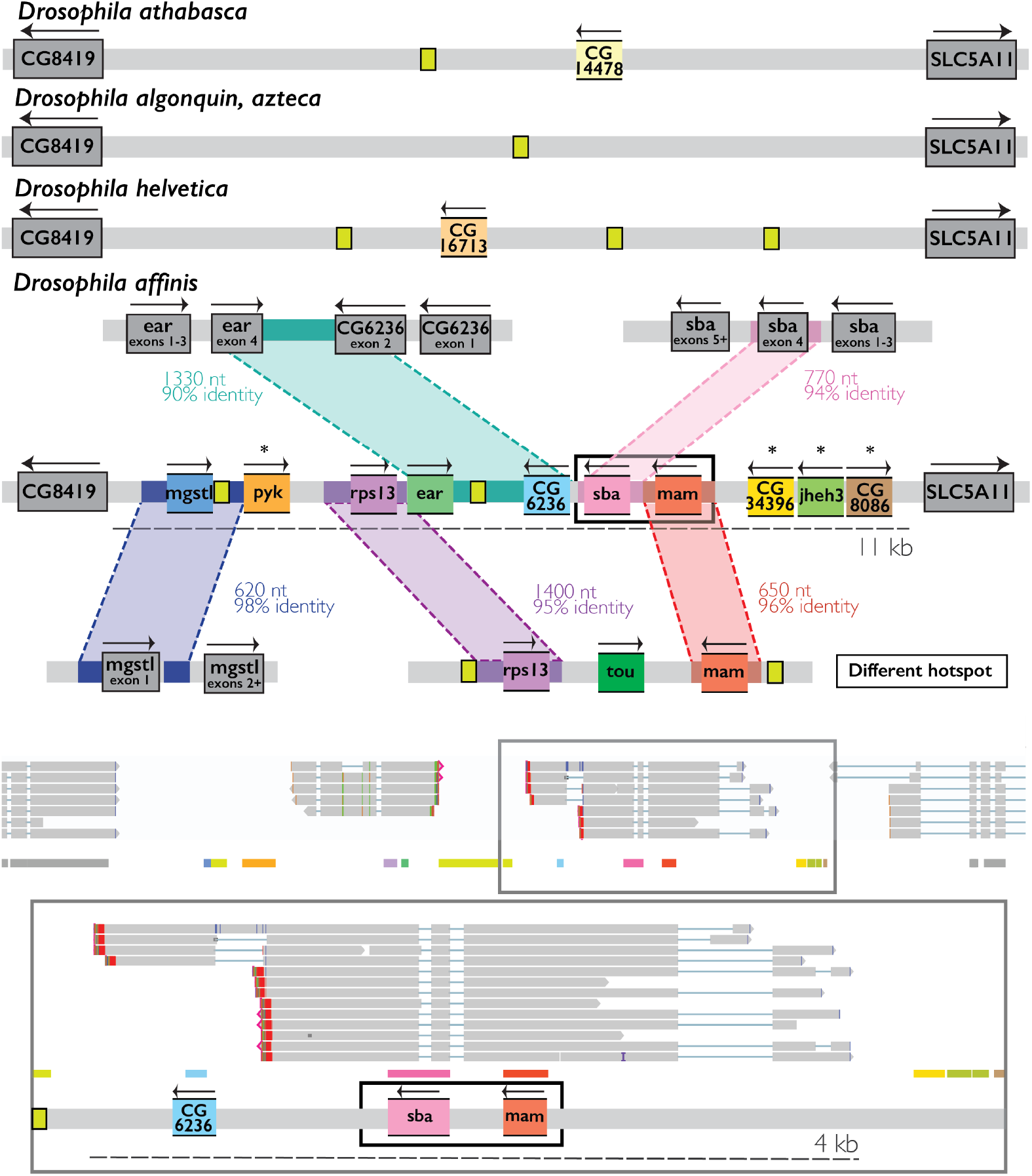
the evolution and transcript structure of a composite ORF made from *mastermind* (mam) and *six-banded* (sba). Details for interpreting the figure are the same as in Figure 1. Top: the orthologous locus in the four closest relatives of *Drosophila affinis*, which do not appear to be hotspots. Middle: the locus in *affinis*, with ten gene fragments, absent in all outgroups. Also shown are the inferred origins via ectopic recombination of the fragments from elsewhere within the *affinis* genome. Starred fragments have no flanking sequence matching noncoding regions in the genome, consistent with ectopic recombination or, less probably, retrotransposition. Black box indicates the composite ORF. Bottom: visualizations of Iso-seq data showing transcripts from the whole locus (top row) and a close-up of those with the *mam/sba* chimeric ORF (bottom row). Thin horizontal colored rectangles directly beneath the reads are the annotations of the gene fragments within IGV; cartoon depictions matching the middle panel are beneath these.

Four of these fragments (*mgst, ear, CG6236, sba*) have flanking sequence with high similarity (90-99%) to intergenic or intronic sequence directly adjacent to exons of their parental genes, strongly suggesting ectopic recombination from there. Two of these (*ear, CG6236)* are exons of neighboring convergent genes and were moved by a single recombination event.

Another four fragments (Figure 2, asterisks) correspond to internal regions of exons. For the same reason as above, exon shuffling seems likeliest, though retrotransposition cannot be excluded.

Based on highly similar (>95%) matching flanking sequence, two fragments (*rps13, mam*) seem to have come not from their ancestral copies, but from another hotspot. Here they are also present in *athabasca*, suggesting that they moved there first from their ancestral loci, and then to the current hotspot. It is not clear whether they were moved together in one recombination event or separately in two.

Two fragments, from homologs of *melanogaster* genes *six-banded* (sba) and *mastermind* (mam), are expressed together on the same transcript at 3 CPM. They form an intact ORF of 576 amino acids, 120 identifiably from *six-banded* and 78 identifiably from *mastermind*.

*Mastermind* is a transcriptional coactivator in the notch pathway important for specifying neural cell fate [14], and *six-banded* is a component of the BAP1 deubiquitination complex, a generalist transcriptional activator [15]. The fragments in the ORF do not correspond to annotated functional domains in either protein (Methods).

There are isoforms of the locus that differ in their 3’ sequence: one contains an additional exon containing a fragment of a third gene *CG8419*. This fragment is in the sense orientation but is not included in the ORF.

Unlike the hotspot in Figure 1, this locus is mostly free of identifiable fragments in all other species: it seems to have become a hotspot only recently. This has coincided with a strong expression increase in a conserved gene flanking the locus, *CG8419* (left, gray in Figure X). In both strains of sister species *athabasca*, and in *Drosophila melanogaster* (FlyAtlas2 [16]), it is expressed at less than 1 CPM. In *affinis*, it is expressed at ∼100 CPM, a 100-fold increase.

#### CSN5/Hph

Ancestrally, the two genes *Hph* and *Tim17a2* are “nested” [17]: the single-exon *Tim17a2* is within the intron of *Hph*, next to and convergently oriented with *Hph’s* first exon (Figure 3). Before the divergence of *affinis* from *athabasca*, a single shuffling event transferred both to a different locus: next to the conserved gene *CSN5*. In *affinis*, there was then substantial duplication and rearrangement of the two gene fragments with their new neighbor *CSN5* at this locus. This created an intact ORF containing sequence from *CSN5* and *Hph*. This ORF was then duplicated within the locus to create a second copy.

**Figure 3:**
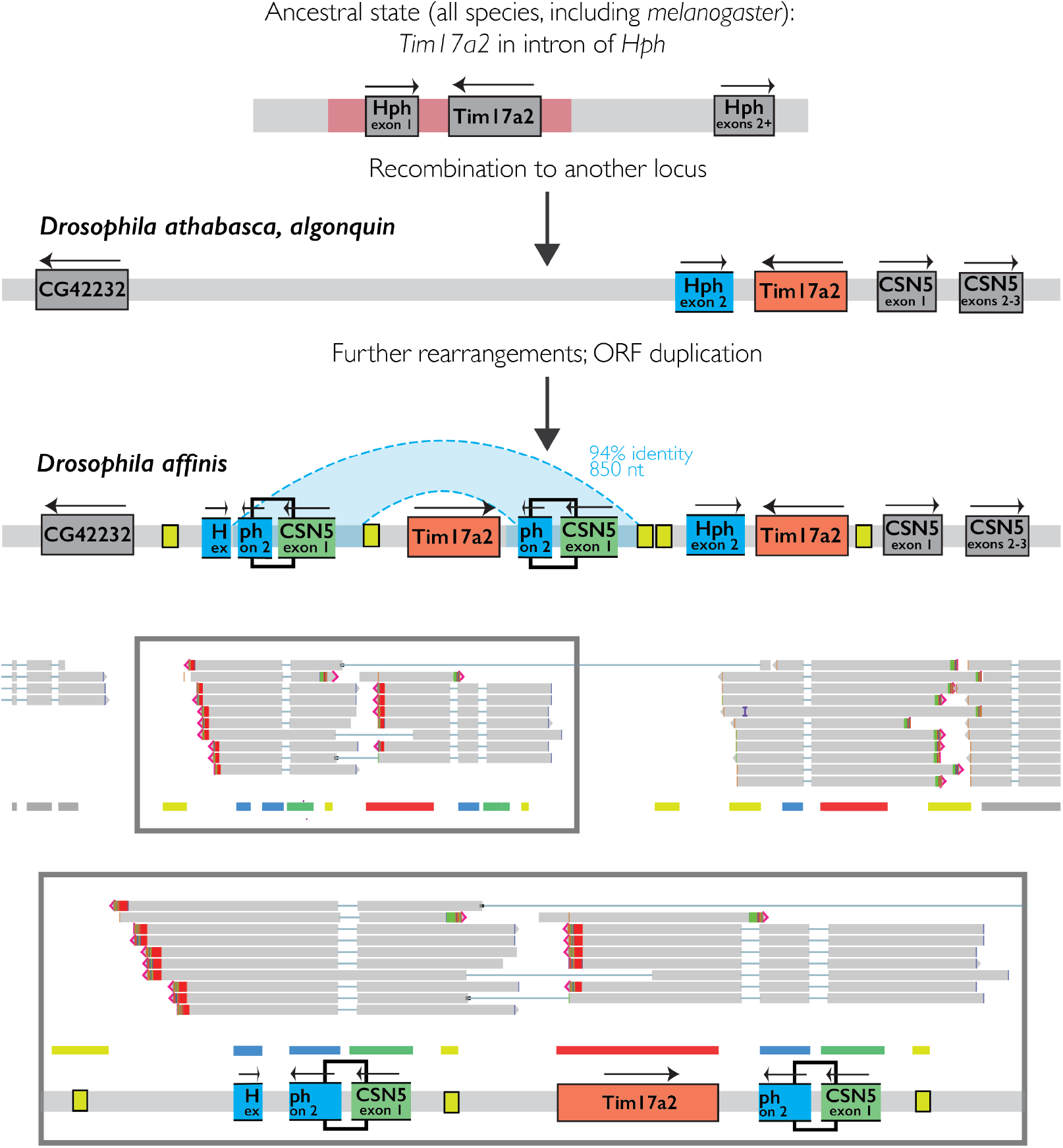
The evolution and transcript structure of a pair of composite ORFs made from *COP9 signalosome unit 5* (*CSN5*) and *Hif prolyl hydroxylase* (Hph). Details for interpreting the figure are the same as for Figure 1. Top: formation mechanism of the ORF. Two exons from nested genes were moved together to a second locus near *CSN5* in the common ancestor of the three indicated species. Specifically in *Drosophila affinis*, that locus was extensively rearranged, creating an intact ORF made from *CSN5* and *Hph*, which was then duplicated. Black boxes indicate the ORF. Bottom: Iso-seq data showing the transcript structure of the whole hotspot (top row) and a close-up of the transcripts containing the new ORFs (bottom row).

Though they contain highly similar ORFs, transcripts from the two duplicates contain different additional sequence elements. One contains a transposon fragment at its 5’ end followed by the ORF; the other contains an additional inverted *Tim17a2* fragment at its 3’ end, which presumably forms a 3’ UTR if the transcript is translated. The two transcripts are each expressed at about 1 CPM.

The hotspot also expresses a third composite transcript with no composite ORF, made from only the *Tim17a2/Hph* fragment, in the direction of the *Hph* fragment. The *Tim17a2* fragment, as in one of the transcripts with the new composite ORF, is on the opposite strand at the 3’ end.

The composite ORF is short, only 41 amino acids, with just 13 from *CSN5. CSN5* is a metalloprotease in the COP9 signalosome complex, responsible for removing a ubiquitin-like protein (NEDD8) to regulate various processes including oogenesis and neurogenesis [18].

Though a physically modest contribution, these 13 residues are functionally key, corresponding to the metalloprotease active site (the “JAMM motif” [19]). *Hph* is a hydroxylase that covalently modifies *sima* (*HIF1a*), the main transcription factor in hypoxia response [20]. The residues in the new ORF from *Hph* do not correspond to any identified domains.

#### CG13876/Prp4k

The pieces for a third composite gene were also taken from a pair of neighboring convergently oriented genes, *CG13876* and *Prp4k*. In *affinis*, a single event shuffled the full sequences of both genes to a different locus, adjacent to a transposon-dense region. There they were extensively duplicated and rearranged, resulting a complex 60 kb array containing more than 40 copies of different fragments derived from them (Figure 4). This array contains seven pairs of closely juxtaposed *CG13876* and *Prp4k* fragments that produce transcripts with a composite ORF.

**Figure 4:**
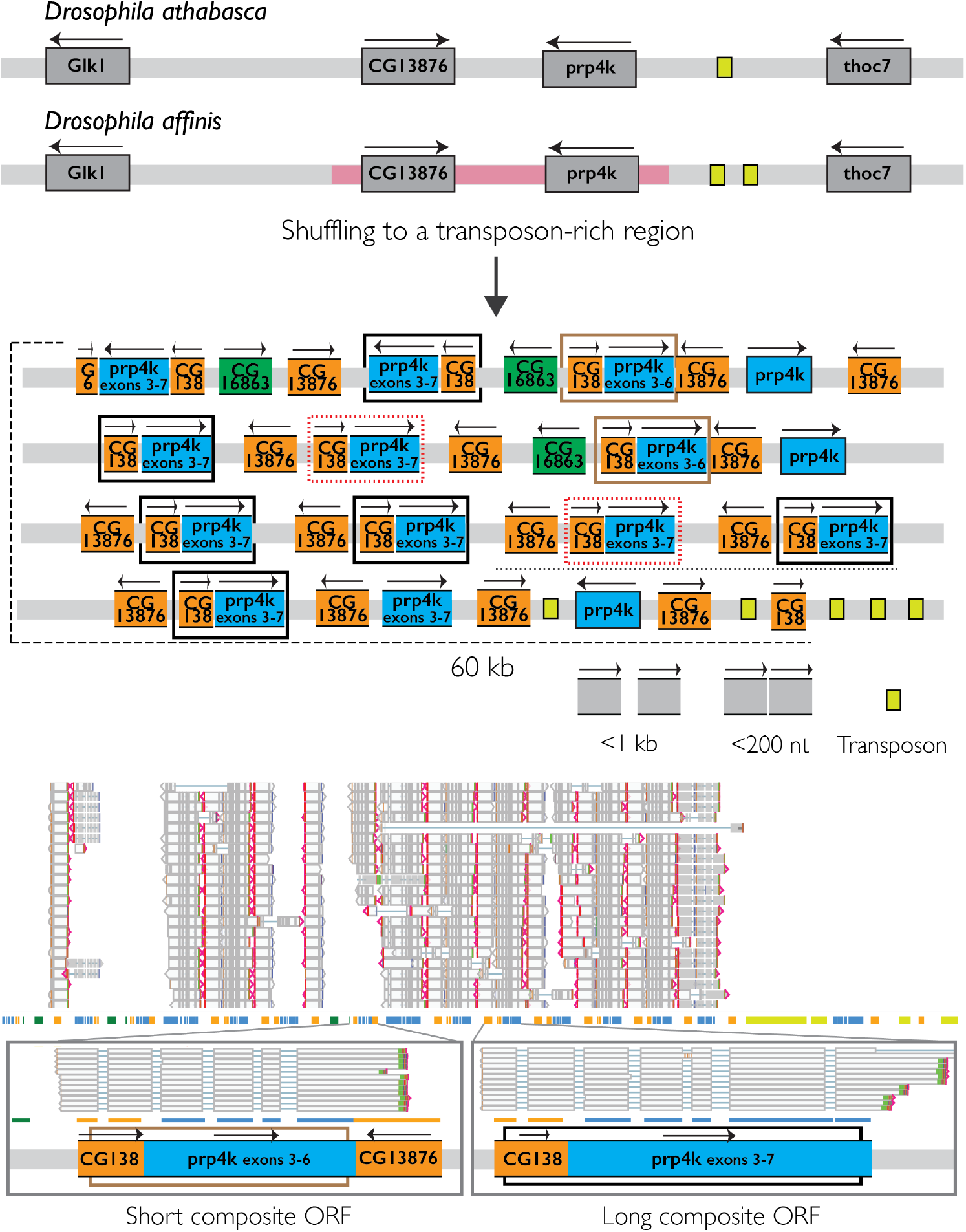
The evolution and structure of an array containing seven copies of composite ORFs made from *CG13876* and pre-mRNA processing factor 4 kinase (*Prp4k)*. Details of interpreting the figure are the same as for Figure 1. Top: cartoon of the locus and its mechanism of formation. Boxes around fragments in the cartoon indicate three types of ORFs: brown boxes are a longer form; black boxes are a shorter form; red boxes are copies that resemble the long form genomically but do not produce an intact ORF due to a truncated 5’ end. Bottom: view of transcripts from the entire locus (top row) and a zoomed-in view of the transcript structure of the long and short forms of the composite ORF (bottom row).

There are two forms of the composite ORF. Five copies in the array have a long form that includes the last exon of *Prp4k*. Two copies have a shorter form, which excludes it and part of the preceding exon, truncating the ORF. This truncation in the short form is due to interruption by a fragment of *CG13874*, longer than the one contained in the ORF, and inverted. If the transcripts are translated, it presumably forms a 3’ UTR. The two forms also have different sequences within 500 nt of the transcriptional state site: the long form is preceded by yet another inverted fragment of *CG13864*, and the long form is preceded by a fragment of a third unrelated gene, *CG16863* (green).

The short forms are each expressed at an average of 1 CPM, for a total of 2 CPM, and the long forms at an average of 6 CPM, for a total of 30 CPM.

The shorter ORF encodes 468 amino acids, 107 from *CG13876* and 341 from *Prp4k*.

The longer form encodes 577 amino acids, 134 from *CG13876* and 473 from *Prp4k. CG13876* is a conserved gene, present in humans, but does not have a known function, and has only one identifiable “domain of unknown function,” comprising the whole 290 amino acid gene. *Prp4k* (*Pre-mRNA processing factor 4 kinase*) is a serine/threonine kinase involved in hippo signaling [21]. Its kinase domain is included in full in both composite ORFs.

The array produces many transcripts other than those described above, corresponding to fragments and combinations of fragments of its components. These include nearly full-length copies of the parent genes; standalone copies of fragments from each of them; the third gene *CG16863*, both on its own and spliced to other elements; and rare splice isoforms linking distant members of the array.

Seeing that these hotspots can rapidly produce new composite coding sequences, I sought a more general sense of their phenomenology.

### Hotspots do not *mostly* create young composite open reading frames

I focused above on transcripts encoding a composite protein because their potential for function is easily assessed, but these are not the main product of hotspots. Most composite transcripts do not contain a composite ORF because their fragments are on different strands, in a different reading frame, or have an intervening stop codon.

In total, I found 226 total composite transcripts from 105 loci (some hotspots produce multiple transcripts, as in Figures 2, 3, and 4 above) that could not be detected in any of outgroups *Drosophila obscura, Drosophila pseudoobscura*, and *Drosophila melanogaster*. Many likely evolved within the *affinis* subgroup. It is possible that some fragments are older and have merely diverged beyond the limits of similarity detection, but the very short evolutionary timescales involved here make this relatively unlikely [22]. At the shortest timescale, I find 166 composite transcripts from 64 loci that appear specific to *Drosophila affinis*.

The formation mechanisms of these composite transcripts do not appear different from their coding counterparts. Two *affinis*-specific examples are shown below in less detail than examples above (Figure 5). In one case (top), a gene was copied in full next to a second gene and pieces of both were mixed together to create a complex, which was then duplicated to form an array of composite transcripts. In another case (bottom), three gene fragments were moved to a relatively small region and formed two convergent transcripts, meeting in the middle of one of the genes and splitting it in half.

**Figure 5:**
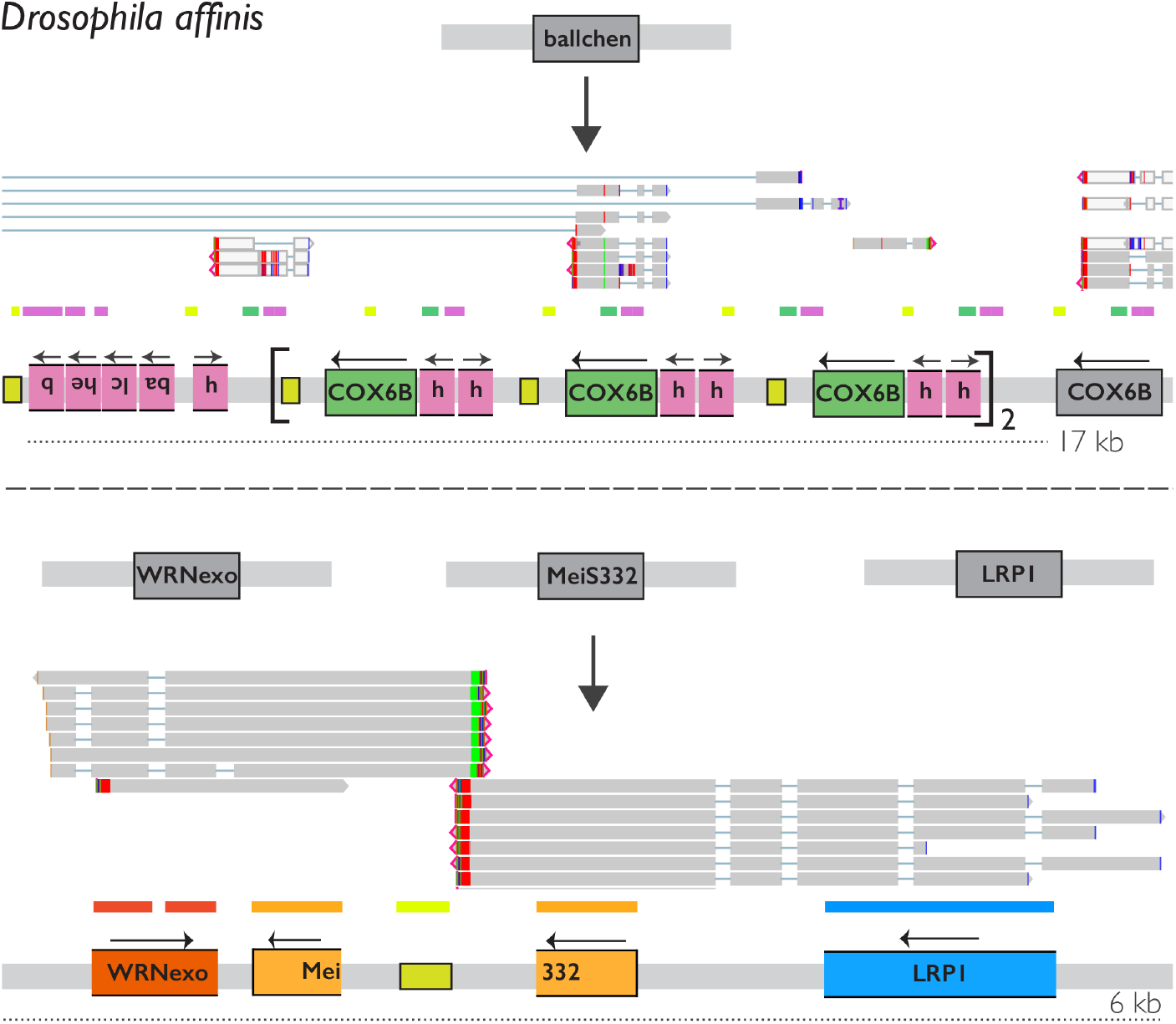
Two hotspots with products other than composite ORFs. Details for interpreting the figure are the same as for Figure 1 above. Top: formation of a composite made from *COX6B and Ballchen. Ballchen* was copied to the *COX6B* locus, where it was disassembled (left), one small fragment (∼50 aa) copied, put head-to-head, juxtaposed with *COX6B*, and duplicated to form an array, among which there is significant diversity in splice isoforms. Bottom: formation of two directly adjacent composite transcripts made from three genes *WRNexo, MeisS332*, and *LRP1. MeiS332* has been split into two pieces, each of which is included along with one other fragment on two transcripts resulting from the locus.

Hotspots produce two other sorts of products: “singletons,” transcripts containing only one identifiable gene fragment, including duplicates; and new exons of ancestral genes. I do not focus on these here because they are comparatively well-understood processes, but their high density at hotspots is noteworthy.

### Variations in amount and kind of hotspot activity

A hotspot in one of these species is often also a hotspot in another (Figure 1, Figure 3), suggesting that they are determined by some feature of the locus that can persist over time.

They can also emerge: the *six-banded/mastermind* hotspot became active very recently (Figure 2).

There are finer gradations between “on” and “off” states in the activity of the same hotspot across species. In Figure 1, one species (*algonquin*) has what appears to be a comparatively quiet hotspot. Figure 6 below shows a comparison of a locus from the preceding section between *affinis* and two close relatives. It is a hotspot in all three species, but has accumulated more fragments in *athabasca*, and many more in *azteca*.

**Figure 6:**
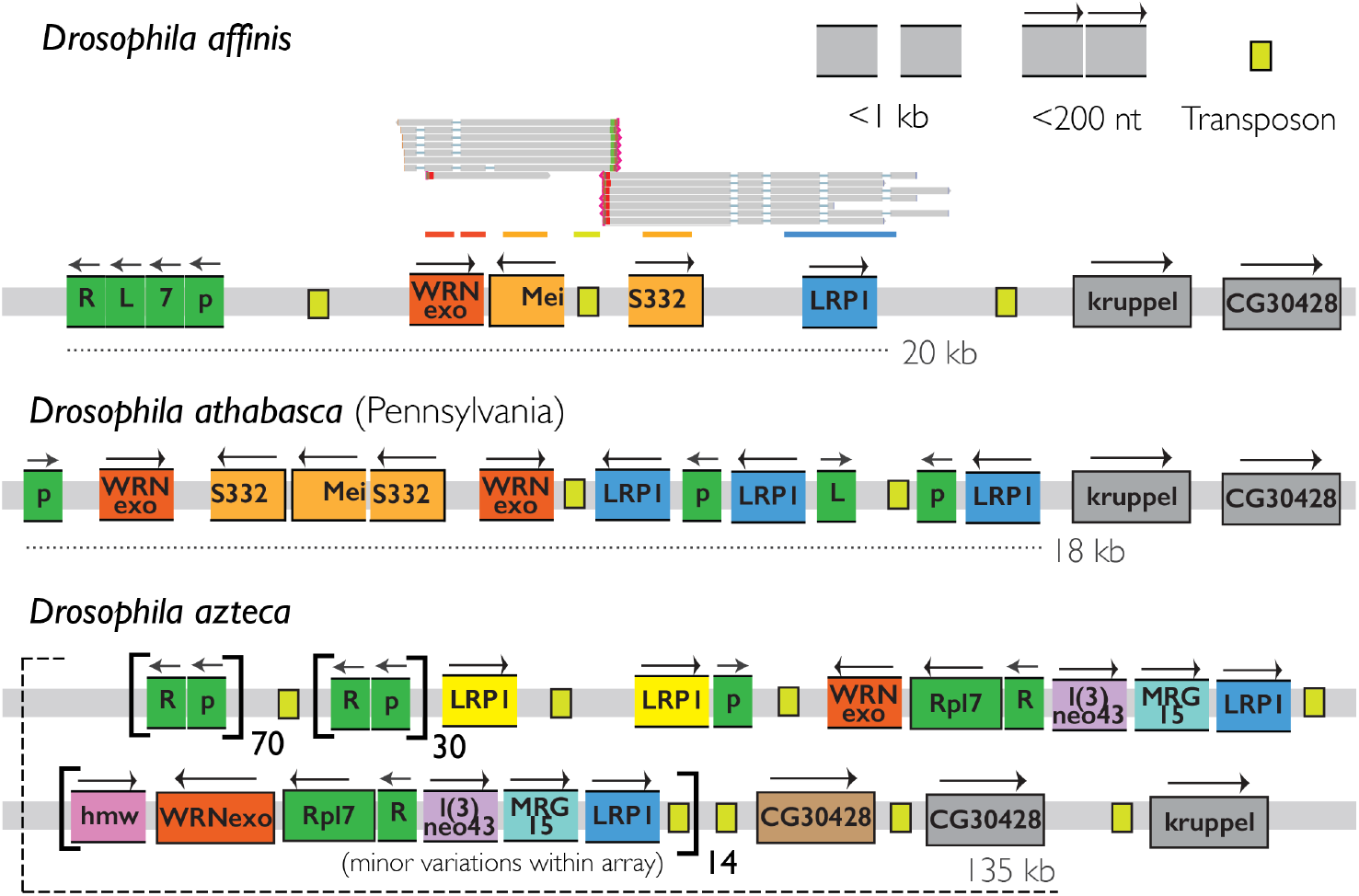
Activity at the same hotspot varies across species. Details of the figure are the same as Figures 1-5 above. Top: a hotspot in *Drosophila affinis*, which gives rise to a transcript lacking a composite ORF. Middle: the same hotspot in *Drosophila athabasca*. The identity of the fragments are the same as in *affinis*, but are duplicated and rearranged. Bottom: the same hotspot in *Drosophila azteca*. Many fragments from the other two species are present, but several new ones have been recruited.Large arrays of extensively rearranged and duplicated fragments have formed in this species.

Making a parsimony-based assumption that the ancestral fragment set includes those present in all species, this increased activity seems to be partitioned into two sorts. In *athabasca*, the higher number of fragments at this locus is due to the duplication and rearrangement of the ancestral fragment set; in *azteca*, it is also due to the gain of several new fragments. This is expected with recombination as a mechanism: repeats occurring within the hotspot allow local rearrangements, and repeats at other loci allow new exon shuffling into the hotspot. The formation of arrays could feed forward into further slippage and rearrangements.

### Hotspots as hubs in recombination networks that spread and receive gene fragments

Not all genes are equally likely to spin off fragments to elsewhere in the genome. Some genes have generated many fragments while others have generated none.

An evocative example is *argonaute 2 (ago2)*, a component of the siRNA and antiviral defense pathways [23]: I find at least 60 *ago2* fragments in the *affinis* genome (Methods).

Extensive adaptive duplication of *ago2* in this species group via DNA-based mutational mechanisms has been previously reported [24]. I find isolated fragments in addition to duplicates, and several hotspots rich specifically in fragments from within the piwi domain, which is the catalytic domain for RNAse activity (all *ago2* fragments, light blue, Figure 1).

Other common fragments can also be seen in the examples above. *CG16863*, the fragment 5’ of some transcripts in the large *CG13876/Pmk4* array (Figure 3, dark green), has 30; *ballchen (*Figure 5, pink), a component in a repeating array, is at three additional loci, at one of which it is a member of different multi-component array; *alpha-spectrin*, a member of the five-fragment composite that is transcribed together in Figure 1 (yellow, orange), has five.

The simplest model of how gene fragments spread heritably around the genome is that a fragment is copied to another locus with a probability defined by its degree of “recombinational contact” in the germline with that locus. This is likely a complex function that includes the loci having sequences that are sufficiently similar for recombination, but also other features of the germline DNA damage and repair process, like the rate at which the locus incurs double-strand breaks, how close in 3D space the two loci are, and so on. Once a fragment successfully spreads, it gains access to the recombinational contacts of its new locus, further increasing its probability of spread. This is a “preferential attachment” model, which is known to result in a power law distribution, f(x) ∝ x^α^ [25], here the number of fragments per gene.

After attempting to “deduplicate” fragment counts that result from processes not related to this model, like further in situ breakdown of fragments into multiple pieces at a new locus (e.g. Figures 5a and 6) or simple tandem duplication (e.g. *ago2* fragments in *azteca*, Figure 1, and Figure 4), and removing fragments that may be moving as or with substantial contribution from retrotransposons (Methods), a power law fits this frequency distribution well (Methods), with a K-S statistic (D), the maximum difference between the CDF of the data and the fit, which can range from 0 to 1 [26], of only 0.03, and an alpha of about 2.6, which is common in biological systems and close to the value of 3 predicted by pure preferential attachment [25] (Figure 7). There are two strong outliers to the fit, both with too many fragments, one of which is *argonaute 2*. This likely reflects positive selection on these fragments, consistent with the previous reports of adaptive *ago2* duplication in the group [24]. Though this model is very simple and neglects many factors (for example, not all loci will have the same number of recombinational contacts, discussed below), its good fit and interpretable deviations support it as a reasonable null model for the process of fragment dissemination.

**Figure 7:**
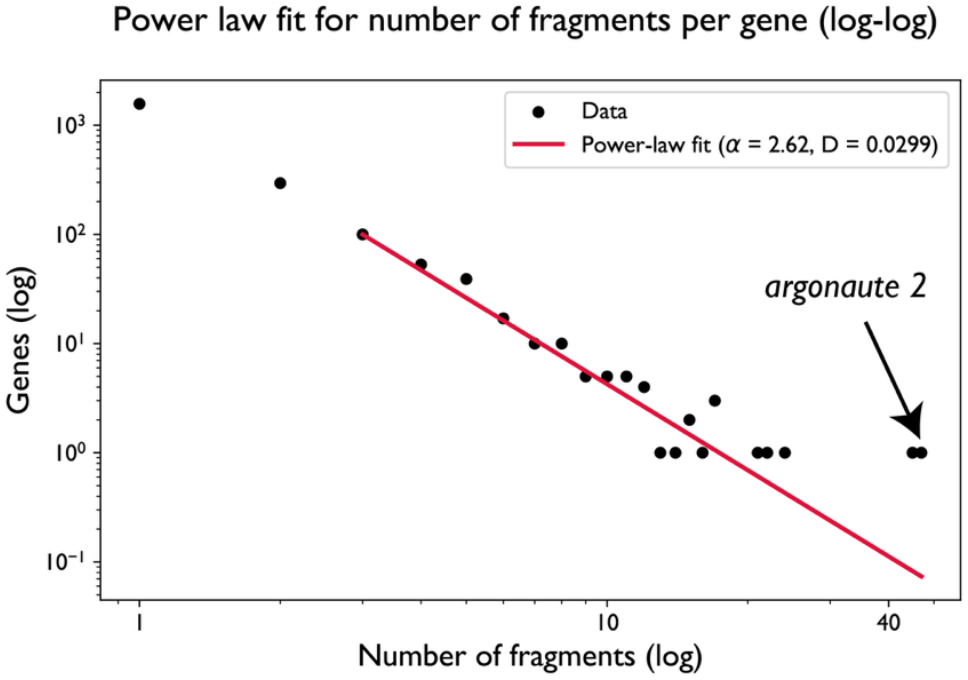
A power law distribution for the number of genomic fragments per gene. x-axis is the number of fragments for a given gene found in the *Drosophila affinis* genome; y axis is the number of genes displaying that number of fragments. Red line is the best fit to a power law distribution, f(x) ∝ x^-α^. Because real distributions often follow a power law at higher values but deviate at low counts, it is standard in power law fits to estimate a lower bound x_min_ at which the power law begins; the red fit line only extends to this value of x_min_=3. *Argonaute 2* is a strong outlier, suggestive of selection operating atop a background process of the spread of gene fragments by recombinational networks.

While they do not appear as non-neutral as *ago2* in this network analysis, some of these less-widely disseminated fragments display interesting properties. Fragments from *Hph* and *Tim17a2*, the components of the complex that “seeded” the *CSN5/Hph* composite ORF (Figure 3), number 10 and 6 respectively, 3 of which are together as a pair in the same head-to-head complex that seeded the ORF. This complex appears in several copies in the genomes of *azteca, athabasca*, and *affinis*, suggesting that it emerged and spread before their common ancestor. Two copies in *affinis* and their transcript structures are shown below. These copies show interesting context-dependent transcription. In an isolated genomic context (Figure 8, top), this complex is transcribed in the direction of one fragment. When placed 5’ of a different gene (Figure 8, bottom), the first fragment is still transcribed, but, in addition, the other fragment is included as a noncoding 5’ UTR for the neighboring gene.

**Figure 8:**
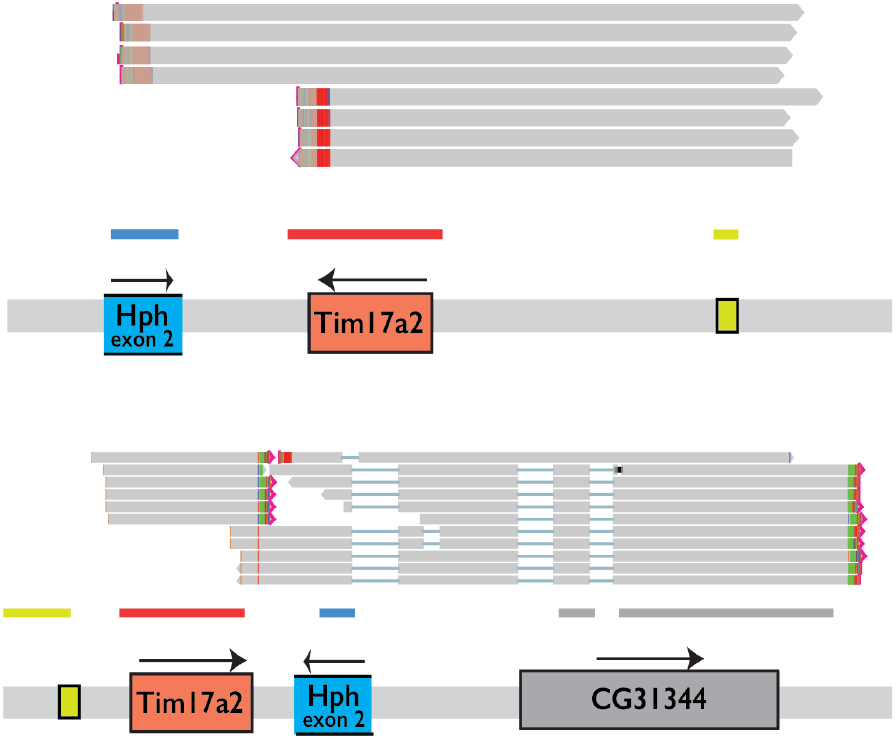
Context-dependent transcription of a common gene fragment. Details of the figure are the same as Figures 1-6 above. This *Tima17a2/Hph* composite unit appears several times in the *Drosophila affinis* genome in different contexts, transcribed to produce different products in each.

A network framework suggests one cause of hotspots: that they are in recombinational contact with unusually many loci. The number of other loci with which a region shares “recombination-competent” sequences should correlate with total recombinational contacts. With reasonable parameters for recombination-competent sequence based on empirical studies of ectopic recombination (shared regions of >200 nt [27] and >90% identity [28] within 1 kb of the locus [29]), I find that the two most active loci (Figures 1 and 2) are in the top 10%, of recombinational contacts, following a class of hotspots that are much larger than average and may be unfairly advantaged in such an analysis. This result is somewhat sensitive to the above parameters, so I caution against overinterpretation and only conclude that recombinational contact is likely an important factor.

As expected, I observe that hotspots can disseminate, as well as receive, gene fragments, including ones that they have “processed” into different forms, like composites. One example of inter-hotspot transfer is above: the secondhand gain of two of the fragments at the *Mastermind/six-banded* ORF (Figure 1). Another example explicitly showing transfer not just of fragments gained at hotspots but of their processed products is shown below for a three-fragment complex within the five-fragment complex seen previously in Figure 1 (top right) between two loci (Figure 9). As both loci are specific to *affinis*, the direction of transfer is less clear.

**Figure 9:**
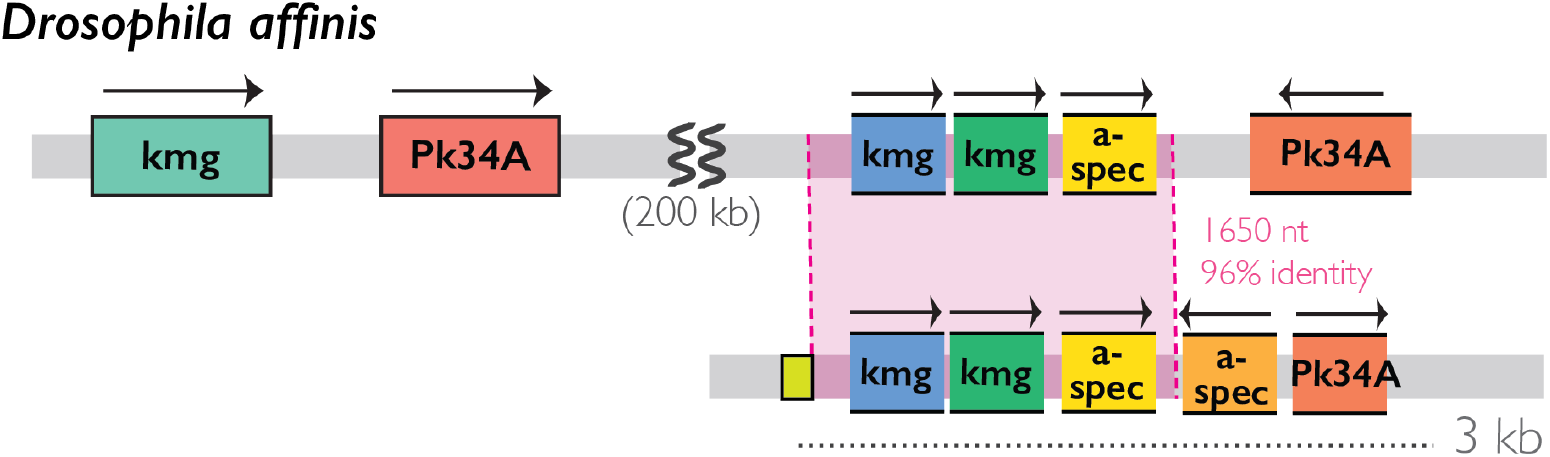
Transfer of a gene fragment complex in *Drosophila affinis*. Details for interpreting the figure are the same as Figure 1 above. The set of five fragments in the bottom right is the same one as shown in Figure 1, top inset. Another locus contains a near-identical copy of three of these fragments, suggesting that this composite has been moved between them. The direction of transfer is not clear.

## Discussion: “Tinkering loci” and the evolutionary reassortment of modular sequences

In the main text I refer to these regions as “exon shuffling hotspots,” but I do not think that this is a particularly good description. For one thing, as observed for other chimeric genes [5], the fragments at these hotspots are not particularly obedient of exon boundaries; sometimes they are several exons, and sometimes they are just pieces of one. For another, “hotspot” to me connotes a high rate of a specific event, but one of the most striking things about these regions is the variety of different structures that they quickly produce, and how surprisingly elaborate those structures can be. I have emphasized visual depictions of these regions to convey this point.

Despite this diversity, I do see these regions as a kind. They are loci that receive sequence fragments from elsewhere around the genome to concentrate and rearrange them. I refer to them by this general tendency to toy [30] with sequences: as “tinkering loci.” This is an homage to François Jacob, who influentially articulated the idea of evolution as a “tinkerer,” putting together pieces that it finds lying around rather than inventing anew [31]. Tinkering loci are workbenches of tinkering in the genome.

We have seen evocative cases of tinkering. Disparate fragments nestled just a few hundred nucleotides apart from one another (Figure 1). Two genes fragmented, as though disassembled, into many small pieces, one carried as a component into a larger structure then duplicated into many copies (Figures 5,6). A gene split in half, its midpoint where two convergent transcripts, with two other fragments, meet and terminate (Figure 5). A single dyadic element, one full gene plus the exon of another, dispersed around the genome, copies contributing independently to new composite elements (Figures 3, 8) and expressed on their own (Figure 8). Two genes unfurled, with a small contribution from a third, into an array structure that looks like a new gene family: a set of well-expressed transcripts, encoding different variants of a new composite protein, with different 3’ UTRs, sitting behind different sequences, conceivably regulatory, just upstream of their TSS (Figure 4). The structures and mechanisms of tinkering loci have the outward trappings of much biological complexity.

Tinkering loci seem like a good solution to the problem, raised in the Introduction, that stands between the idea of exon shuffling and its maximum potential as an engine that reassorts functional sequences that evolution has on hand: the improbability of those sequences bumping into one another in just the right way in the vast animal genome. They are an even better solution if their features also enable them to copy and spread successful products. Most of their products are likely not useful, but with time and numbers, they can hit upon a few that are.

It is not currently clear whether tinkering loci are general to animals or a peculiarity of the *affinis* group. A related question, essential but unanswered, is what exactly causes tinkering loci. It seems likely that the *affinis* group has more tinkering activity than average, based on having encountered them here and on previous reports of unusually many apparently novel genes in the group [32,33]. It is interesting to speculate whether tinkering loci could underlie exotic phenomena found in these flies, like sex ratio meiotic drive [34] and heterospermy [35].

Similarly, it is tempting to make stories of how, if they are general, tinkering loci could help to explain and organize animal evolution. Saltational processes could have a mechanistic basis in the changing activity of tinkering loci over time, perhaps induced by an influx of recombination substrates in the form of transposons. I have focused here on protein elements, but it is possible that regulatory elements are also at tinkering loci, which would then appeal as a mechanism for the complex regulatory grammar that evolved quickly in animals. The word grammar here begins to feel literal: one can see faintly above the suggestion of characteristic patterns of genomic motifs emerging from how tinkering loci tend to recurrently apply their mutational functions of insertion, deletion, duplication and rearrangement. The repeat structure within a tinkering locus, or the recombinational network between them, could influence and explain what sorts of composites tend to emerge.

## Methods

### Iso-seq of male *Drosophila affinis*

*Drosophila affinis* strain 14012-0141.00 was purchased from the National Drosophila Species Stock Center (NDSSC). This is the strain used to generate the reference assembly.

Approximately 40 male flies, ages 3-5 days, were anesthetized with CO_2_, frozen on dry ice, and homogenized in Zymo DNA/RNAShield. RNA was extracted using the ZymoBIOMICS Quick-RNA miniprep kit (Zymo Research, R1655) and eluted into 30 uL. Iso-seq libraries were created according to the ‘Preparing Iso-Seq® v2 libraries’ protocol from PacBio, with barcoded cDNA primers for multiplexing with three other *Drosophila* samples. Libraries were pooled, bound, and cleaned according to the Revio SPRQ™ Polymerase kit. Libraries were then loaded onto the Revio instrument and run using the application presets.

### Genome annotations

To annotate protein-coding genes and fragments, I performed a tBLASTn search [36] of each genome assembly using the RefSeq protein set for *Drosophila melanogaster* (Resources and Data Availability) as query with an E-value threshold of 10^-3^. This approach considers proteins in the high-quality *melanogaster* annotation as a rough proxy for the ancestral protein set from which fragments may be derived. It has the drawback that fragments generated directly by proteins in *affinis* group species that had already diverged too far to be detected by their *melanogaster* homologs will be missed.

I then partitioned the results of this BLAST search into orthologs of the *melanogaster* genes and duplicates, including fragments, relative to *melanogaster*. For each *melanogaster* gene, I identified the best candidate ortholog by finding the hit with the highest statistical significance (lowest E-value) that also includes at least 60% of the protein (potentially separated by introns of up to 50 kb). This process identified orthologs for 97% of *melanogaster* genes including an average of 85% of their length (not removing 0 values from missing orthologs). Though this process is somewhat heuristic, I note that conserved genes would be filtered out of the final list of new composite transcripts in the evolutionary analysis.

I then identified every BLAST hit that did not overlap with these orthologs and estimated its boundaries and closest *melanogaster* relative by selecting the hit with the lowest E-value. These were my candidate gene duplicates and fragments. I made a second pass through these to identify and condense separate hits that were spuriously identified as from separate proteins by slight stochasticity in the E-values of many related *melanogaster* paralogs. To do this, I made a clustered representation of the *melanogaster* proteome via a BLASTP search with an E-value of 10^-3^, and merged hits in the same cluster and on the same strand within 20 kb of one another. I considered the result from this stage the fragment repertoire of the genomes.

I also annotated transposable elements. I took three approaches, hoping to identify fast-evolving, lineage-specific transposon families. First, I downloaded 881 known transposon sequences from Dfam [37] classified as Dipteran (Resources and Data Availability) used them as a query in a blastn search against each assembly, using an E-value threshold of 10^-3^ to annotate transposon sequences in the genome. Second, I performed a six-frame translation of each assembly using the easel package in HMMER [38] and used it as the target in an hmmscan search with a custom database containing common transposon profile HMMs as a query (Resources and Data Availability), using an E-value threshold of 10^-4^ to annotate transposon-derived genomic regions. Finally, I performed the six-frame translation approach specifically on Iso-seq transcripts that did not match any of my annotated features, reasoning that transposons are comparatively likely to be expressed, which allows a reduction in target size and higher power. I considered unannotated transcripts with hits to these transposon-associated protein domains putative transposons, and annotated them in the *affinis* genome using the same blastn search as for the Dfam sequences.

### Identification of composite transcripts

I mapped the Iso-seq reads to the *affinis* genome using minimap2 with parameters -x splice-hq, --splice-flank no, --secondary no. I used my annotation and this mapping to identify reads that contain protein fragments from two or more different *melanogaster* genes.

### Transcript quantification

Iso-seq data for *affinis* were quantified as counts per million (CPM): the number of reads for that transcript divided by the total number of reads in the dataset.

### Evolutionary analyses

I extracted the *affinis* genome sequences corresponding to the full span covered by each composite transcript, including introns. I used these as a query in a dc-megablast search using an E-value of 10^-2^ to search the assembly of each other species for these fragments and any intervening sequence. I preliminarily consider a composite transcript absent in an outgroup species if the genomic sequences corresponding to the fragments are not found within 10 kb of one another in the outgroup genome. Because of the small evolutionary distances involved, identifying the orthologous locus itself was successful in all but one case in outgroup *Drosophila pseudoobscura* even when the proteins there could not be detected, making significant divergence beyond detection unlikely [22]. For cases presented here, I also searched each species’ protein annotation for fragments of the same genes identified in proximity to one another. I note that it is possible that the genome annotation process above conflates occasional pieces of ancestral genes with new fragments by its own lights, but that this procedure means that these ancestral “fragments” would be excluded from the final set of chimeric transcripts due to being conserved in *affinis* group species.

### Protein domain identification

All protein domain analysis were performed with both hmmscan at https://www.ebi.ac.uk/Tools/hmmer/search/hmmscan using the Pfam 37.2 release and with the C-D search tool on the BLAST web server (https://blast.ncbi.nlm.nih.gov/Blast.cgi).

### Power law analysis

To perform the power law fit (D statistic) and calculate the best fit value for alpha, I used the powerlaw package for Python [39].

To deduplicate raw counts of protein fragments (Genome annotation) into counts more representative of the spread of fragment by ectopic recombination, I merged fragments of the same gene within 10 kb of one another on the same strand with no other sequences intervening. This is intended to eliminate multiple counts resulting from in situ fragmentation of the gene into multiple apparent fragments, e.g. by insertions or deletions within the gene, the beginnings of sequence degradation. It is also intended to eliminate simple tandem duplications.

I also removed all protein fragments that frequently overlap with genome regions that my transposon annotations have also called as lineage-specific transposons (Genome annotations). The motivation for this was that these proteins may be transposons or frequently moving by retrotransposition, another process that does not reflect the model being assessed here.

## Resources and data availability

### Genome assemblies

*<u>Drosophila affinis</u>*:This assembly is from a population collected in Hastings, Montana, in 1950. A preliminary version of an assembly generously shared by Bernard Kim was used for this work. It is very similar to GCA_035045985.1 in Genbank, the final polished version presented in [12]. The version used here is available at https://caraweisman.com/data.

*<u>Drosophila athabasca</u>* (New Jersey): this assembly is from an isofemale line, NJ28, collected in Princeton, New Jersey in 2016 and presented in [13]. It is available in GenBank as GCA_008121215.1.

*<u>Drosophila athabasca</u>* (Pennsylvania): this assembly is from an isofemale line, PA60, collected in Black Moshannon, Pennsylvania, in 2010, and presented in [13]. It is available in GenBank as GCA_008121225.1.

*<u>Drosophila algonquin:</u>*<u>this assembly is from an individual collected in the Huron Mountains, Michigan, in 2021</u>. A preliminary version of an assembly generously shared by Bernard Kim was used for this work. It is very similar to GCA_035041765.1 in Genbank, the final polished version presented in [12]. The version used here is available at https://caraweisman.com/data.

*<u>Drosophila azteca</u>*: this assembly is from strain UCSD 14012-1071.03 and is available as GCA_005876895.1 in Refseq.

*<u>Drosophila helvetica</u>*<u>(annotation used in main text):</u> This assembly is from an individual collected in 2021 in East Sussex, England. A preliminary version of an assembly generxously shared by Bernard Kim was used for this work. It is very similar to GCA_037043705.1 in Genbank, the final polished version, which was presented in [12]. The version used here is available at https://caraweisman.com/data.

*<u>Drosophila helvetica</u>*<u>(Figure 1):</u> Figure 1 shows the primary and alternate assemblies resulting from a different sequencing project for *helvetica*, only used for this figure. These are available in GenBank as GCA_963969585.1 and GCA_963969575.1 and were presented in [11].

*<u>Drosophila obscura</u>*<u>: this assembly is available as</u> GCA_018151105.1 in GenBank.

*<u>Drosophila pseudoobscura</u>*: this assembly is available as GCA_009870125.2 in GenBank.

*<u>Drosophila melanogaster</u>*: this is the standard reference assembly, available as GCF_000001215.4 in RefSeq.

All of these assemblies are based on long read sequencing technologies (Pacbio or Oxford Nanopore).

### RNA sequence data

*<u>Drosophila athabasca (New Jersey)</u>*:I used Illumina short read RNAseq available from the testes of males of the “EB” strain of *athabasca*, available in the SRA as SRR9967660 and presented in [13], which specifies that this is the same as line NJ28, used for the assembly.

*<u>Drosophila athabasca (Pennsylvania)</u>*: I used Illumina short read RNAseq available from the testes of males of the “EA” strain of *athabasca*, available in the SRA as SRR9967656 and presented in [13], which specifies that this is the same as line PA60, used for the assembly.

*<u>Drosophila affinis</u>*: I generated a new Iso-seq dataset from this species according to the Methods above, available at https://caraweisman.com/data.

### Genome annotations, coordinates of composite transcripts, and queries for transposon annotation

Genome annotations coordinates of composite transcripts including all tinkering loci highlighted in this analysis, and files used for annotation of transposons are available at https://caraweisman.com/data.

## Acknowledgements

I am grateful to Joshua Akey for use of laboratory space and equipment; to Joshua Akey, Julien Ayroles, and Michael Levine for mentorship and helpful conversations; to members of the Ayroles and Levine labs for experimental support; to Sean Eddy for computing resources; to Sean Eddy and Andrew Murray for helpful conversations; and to Bernard Kim for his early and generous sharing of many of the genome assemblies used here. I am especially grateful to the Lewis-Sigler Institute for Integrative Genomics at Princeton, the source of all funding.

